# Microbial metabolite deoxycholic acid shapes microbiota against *Campylobacter jejuni* chicken colonization

**DOI:** 10.1101/584284

**Authors:** Bilal Alrubaye, Mussie Abraha, Ayidh Almansour, Mohit Bansal, Hong Wang, Young Min Kwon, Yan Huang, Billy Hargis, Xiaolun Sun

## Abstract

Despite reducing the prevalent foodborne pathogen *Campylobacter jejuni* in chickens decreases campylobacteriosis, few effective approaches are available. The aim of this study was to use microbial metabolic product bile acids to reduce *C. jejuni* chicken colonization. Broiler chicks were fed with deoxycholic acid (DCA), lithocholic acid (LCA), or ursodeoxycholic acid (UDCA). The birds were also transplanted with DCA modulated anaerobes (DCA-Anaero) or aerobes (DCA-Aero). The birds were infected with human clinical isolate *C. jejuni* 81-176 or chicken isolate *C. jejuni* AR101. Notably, *C. jejuni* 81-176 was readily colonized intestinal tract at d16 and reached an almost plateau at d21. Remarkably, DCA excluded *C. jejuni* cecal colonization at 100, 99.997, and 100% at 16, 21, and 28 days of age. Neither chicken ages of infection nor LCA or UDCA altered *C. jejuni* AR101 chicken colonization level, while DCA reduced 91% of the bacterium in chickens at d28. Notably, DCA diet reduced phylum Firmicutes but increased Bacteroidetes compared to infected control birds. Importantly, DCA-Anaero attenuated 93% of *C. jejuni* colonization at d28 compared to control infected birds. In conclusion, DCA shapes microbiota composition against *C. jejuni* colonization in chickens, suggesting a bidirectional interaction between microbiota and microbial metabolites.

## INTRODUCTION

*Campylobacter jejuni* asymptomatically colonizes in poultry gut but is one of the prevalent food borne pathogens in the developed countries. Antibiotic resistant *C. jejuni* has been reported in children and adults in US and worldwide [1–5]. More than 13 campylobacteriosis cases per 100,000 population were recorded in 2014 in USA, which represents a 13% increase compared to 2006-2008 and was higher than the combined incidences by the following 8 bacterial pathogens [6]. A total of 1.3 million individuals are afflicted by the disease, resulting in 76 deaths every year [7]. Although antibiotics treatment has a marginal benefit (1.32 days) on the duration of campylobacteriosis symptoms [8], it is the only current available therapy for patients with severe campylobacteriosis or for those at high risk for severe disease [9]. However, ever increasing antimicrobial resistance [10] prompts the need for immediate and sustainable counter-actions from agricultural industry to medical field. Misuse or overuse of antimicrobial agents in medical and agricultural practice is contributing to exacerbating the episodes of emerging antimicrobial resistant microbes [10]. Hence, an effective and sustainable solution is to find antimicrobial alternatives in agricultural industry and medical field.

Tremendous efforts have been put forward to prevent campylobacteriosis by reducing *C. jejuni* contamination in animal food sources, particularly broiler chicken meat. The intervention approaches include on-farm biosecurity measures [11], vaccines [12], probiotics [13], phages [14], and post-slaughter decontamination of poultry carcasses [15]. Reducing carcass *Campylobacter* counts by 2 log is estimated to decrease a 30-fold in human campylobacteriosis [16]. Although those measures to prevent *C. jejuni* contamination have achieved some success, improvement is needed, as evidenced by the relative consistent rate of campylobacteriosis incidences in the Morbidity and Mortality Weekly Report at CDC infectious disease database from January,1996 to June, 2017 [17].

Sparse information is currently available on using microbiota to prevent *C. jejuni* colonization in poultry. Microbiota transplantation has shown tremendous success against recurrent *Clostridium difficile* infection [18]. The production of secondary bile acids by specific bacteria are attributed to inhibit *C. difficile* colonization and infection [19]. Bile acids at 3-30 mM in the human small intestine [20] are the byproducts of cholesterol and biotransformed from conjugated to unconjugated primary bile acids, and to secondary bile acids. Majority of bile acids (>95%) are effectively absorbed in intestine [21]. Primary bile acids of cholic acid (CA) and chenodeoxycholic acid (CDCA) are synthesized from cholesterol in hepatocytes and conjugated with glycine or taurine [22]. In the intestine, the conjugated primary bile acids are deconjugated by bacterial bile salt hydrolase (BSH) and further altered by microbiota to produce secondary bile acids including lithocholic acid (LCA), deoxycholic acid (DCA), and ursodeoxycholic acid (UDCA). Deconjugating enzyme BSH is present in all major bacterial divisions and archaeal species in the human gut including members of genus *Lactobacilli*, *Bifidobacteria*, *Clostridium*, and *Bacteroides* [21,23–25]. Secondary bile acid producing bacteria consist of a small population of genus *Clostridium*, including *C. scindens*, *C. hiranonis*, *C. hylemonae* (Clostridium cluster XVIa), and *C. sordelli* (Clostridium cluster XI) [21]. Besides emulsification of lipid for digestion, bile acids are implicated in various signaling pathways including nuclear receptors of farnesoid X receptor (FXR) [26], pregnane X receptor (PXR) [27], and vitamin D receptor (VDR) [28], as well as G protein-coupled receptors of G-protein-coupled bile acid receptor (TGR5) [29] and sphingosine-1-phosphate receptor 2 (S1P2) [30]. The bile acid-metabolizing bacteria such as *Lactobacilli* and *Bifidobacteria* are probiotics and they enhance health by promoting host immune homeostasis [31]. Besides bile acids influence host response, their level is associated with microbial community dynamics in the gut [32]. Bile acids directly inhibit gut microbes [33] and indirectly modulate microbiota through FXR-induced antimicrobial peptides [34]. Mice fed CA have increased class Clostridia (70 vs. 39%) compared to control mice, and genus Blautia (including Ruminococcus spp. and Clostridium cluster XIVa) expands from 8.3 to 55-62% [35]. Bile acids, particularly secondary bile acid DCA, are associated with a variety of chronic diseases, such as obesity, diabetes, and colorectal tumorigenesis [36,37]. Only very recently, new evidences shed light on the beneficial property of secondary bile acids in health and diseases, such as gut motility [38] and *C. difficile* infection [19]. We found that specific pathogen free (SPF) *Il10^-/-^* mice resisted against *C. jejuni* 81-176 induced colitis, while the mice were susceptible to campylobacteriosis after treated with antibiotic clindamycin which kills bile acid metabolizing bacteria [39]. 16S rDNA sequencing, bioinformatic, and HPLC/MS analysis showed that clindamycin depleted all secondary bile acids including DCA. Furthermore, anaerobe metabolite DCA prevents and treats *C. jejuni*-induced colitis in ex-Germ Free mice [39]. However, it remains unclear whether DCA regulates *C. jejuni* chicken colonization and transmission.

In this study, we hypothesized that DCA prevented *C. jejuni* chicken colonization. Our data indicate that DCA indeed prevented against chicken colonization of *C. jejuni* human clinical isolate 81-176 or chicken isolate AR101. Subsequent mechanistic studies found that DCA modulated intestinal microbiota and DCA-modulated anaerobes attenuated *C. jejuni* chicken colonization. Thus, the action of DCA against *C jejuni* chicken colonization represents unique bidirectional interaction between microbiota and microbial metabolites. It would be more effective against campylobacteriosis or the pathogen colonization through modulating both microbiota and microbial metabolites.

## MATERIAL AND METHODS

### Chicken experiment

Animal experiments performed were in accordance with the Animal Research: Reporting of *In Vivo* Experiments (https://www.nc3rs.org.uk/arrive-guidelines). The experiments were approved by the Care and Use Committee of the University of Arkansas. Cohorts of 13 to 18 one-day-old broiler chicks obtained from Cobb-Vantress (Siloam Springs, AR) were neck-tagged and randomly assigned to floor pens with a controlled age-appropriate environment. The birds were fed a corn-soybean meal-based starter diet during 0-10 days of age and a grower diet during 11-28 days of age. The basal diet was formulated as described before [40]. Bird were fed diet supplemented with 0 or 1.5 g/kg DCA, LCA, or UDCA (all from Alfa Aesar) from d 0. Before experiment, frozen stock of *C. jejuni* 81-176 and AR101 (isolated at Dr. Billy Hargis’s lab at University of Arkansas at Fayetteville) were cultured for 48 hours on *C. jejuni* selective blood plates with five antibiotics (cefoperazone, cycloheximide, trimethoprim, vancomycin and polymyxin B). Birds were infected with 10^9^ CFU/bird human clinical isolate *C. jejuni* 81-176 at 14 days of age or chicken isolate AR101 at d 5, 10, or 14. Most birds were infected with *C. jejuni* AR101 at day 10 in order to have enough time for both *C. jejuni* colonization and microbiome development. Chicken body weight and feed intake were measured at 0, 14, 21, and 28 days of age. Birds were sacrificed at 16, 21 or 28 days of age to collect cecal samples for enumerating *C. jejuni*. Cecal digesta samples were collected for DNA isolation or were ten-fold serially diluted with sterile PBS and cultured at 42°C for 48 hours under microaerophilic atmosphere. Colonies were enumerated and CFU per gram was calculated.

### Microbiota transplantation and *C. jejuni* colonization

Because *C. jejuni* wasn’t detected in cecal digesta samples from 28 days old birds infected with *C. jejuni* 81-176 and fed 1.5g/kg DCA diet, the samples were used for isolating transplantation microbiota. Briefly, the cecal content from the birds were quickly squeezed into 50 ml conical tubes and PBS and glycerol were added. The suspension was homogenized, aliquoted, and stored in −80 °C freezer. Before experiments, the microbiota stock was cultured on brain heart infusion (BHI) plates under aerobic or anaerobic conditions for 48 hours at 42°C. The collected bacteria were labelled as DCA modulated aerobes (DCA-Aero) and DCA modulated Anaerobes (DCA-Anaero). Chickens were orally gavaged once with 10^8^ CFU/bird DCA-Aero or DCA-Anaero at 0 days of age. At 10 days of age, the birds were infected with 10^9^ CFU/bird chicken isolate *C. jejuni* AR101. Cecal digesta collected at days 21 and 28 were serially diluted and cultured for *C. jejuni* enumeration as described above.

### *In vitro C. jejuni* growth with various bile acids

The impact of various species of bile acids on *C. jejuni* growth was measured. Briefly, frozen *C. jejuni* 81-176 or AR101 were cultured on the select plates for 48 hours. *C. jejuni* at 10^3^ CFU was inoculated into 5 ml Campylobacter Enrichment Broth (Neogen Food Safety, MI) in the presence of DCA, taurocholic acid (TCA), or CA at various concentrations (0, 5, 25 mM) with triplication. The bacteria were cultured for 24 hours at 42 °C under microaerobic condition. *C. jejuni* growth was measured by serial dilution and plating on the select plates.

### Microbiota composition at phylum level

Cecal digesta samples were collected and DNA was extracted using bead beater disruption and phenol: chloroform separation as describe before [39]. The levels of five phylum bacteria were determined using SYBR Green PCR Master mix (Bio-Rad) on a Bio-Rad 384-well Real-Time PCR System. The PCR reactions were performed according to the manufacturer’s recommendation. The following gene primers [39,41] were used: Universal 16S: 16s357F: 5’-CTCCTACGGGGAGGCAGCAA-3’, 16s1392R: 5’-ACGGGCGGTGTGTRC-3’; α-proteobacteria: α682F 5’-CIAGTGTAGAGGTGAAATT-3’, 908αR 5’-CCCCGTCAATTCCTTTGAGTT-3’; γ-proteobacteria: 1080γF 5’-TCGTCAGCTCGTGTYGTGA-3’ γ1202R 5’-CGTAAGGGCCATGATG-3’; Bacteroidetes: 798cfbF 5’-5’-CRAACAGGATTAGATACCCT-3’ cfb967R 5’-GGTAAGGTTCCTCGCGTAT-3’; Firmicutes: 928F-Firm 5’-TGAAACTYAAAGGAATTGACG-3’ 1040 Firm R 5’-ACCATGCACCACCTGTC-3’; Actinobacteria: Act920F3 5’-TACGGCCGCAAGGCTA-3’ Act1200R 5’-TCRTCCCCACCTTCCTCCG-3’. The relative fold change of each phylum in one sample were normalized to universal 16S. The percentage of each phylum was then calculated as the phylum relative folds divided by total folds of all five phyla.

### Statistical Analysis

Values are shown as mean ± standard error of the mean as indicated. Differences between groups were analyzed using the nonparametric Mann–Whitney *U* test or unpaired t-test with Welch’s correction using Prism 7.0 software similar to previous reports [39,42], because some data were not under normality. Experiments were considered statistically significant if *P* values were <0.05.

## RESULTS

### DCA prevents *C. jejuni* strain 81-176 cecal colonization in chickens

Secondary bile acid DCA prevents and treats *C. jejuni*-induced intestinal inflammation in germ free *Il10^-/-^* mice [39]. Because chickens are the natural reservoir of *C. jejuni*, we then interrogated the hypothesis that DCA modulated *C. jejuni* colonization in chickens. The birds fed diet supplemented with 1.5 g/kg DCA were orally infected with a single dose of 10^9^ CFU/bird human clinical isolate *C. jejuni* strain 81-176 at 14 days of age. *C. jejuni* colonization level was determined by collecting and culturing cecal digesta of the birds at 16, 21, and 28 days of age using *C. jejuni* select medium. Notably, no *C. jejuni* in cecal digesta was detected from birds without the bacterial infection, suggesting the clean facility at our chicken farm and the success of strict biosecurity measurement during our experiments. *C. jejuni* was readily colonized the intestinal tract at a level of 10^5^ CFU/g cecal digesta at 16 days of age, only 2 days post infection (Figure 1). *C. jejuni* colonization level then increased more than 100 folds and reached an almost plateau of 2.8 x10^7^ CFU/g cecal digesta at 21 days of age. Remarkably, DCA excluded *C. jejuni* cecal colonization at 100, 99.997, and 100% at 16, 21, and 28 days of age compared to the infected control birds. These results suggest that the secondary bile acid DCA effectively reduces *C. jejuni* 81-176 colonization in the intestinal tract of broiler chickens.

**Figure 1.**
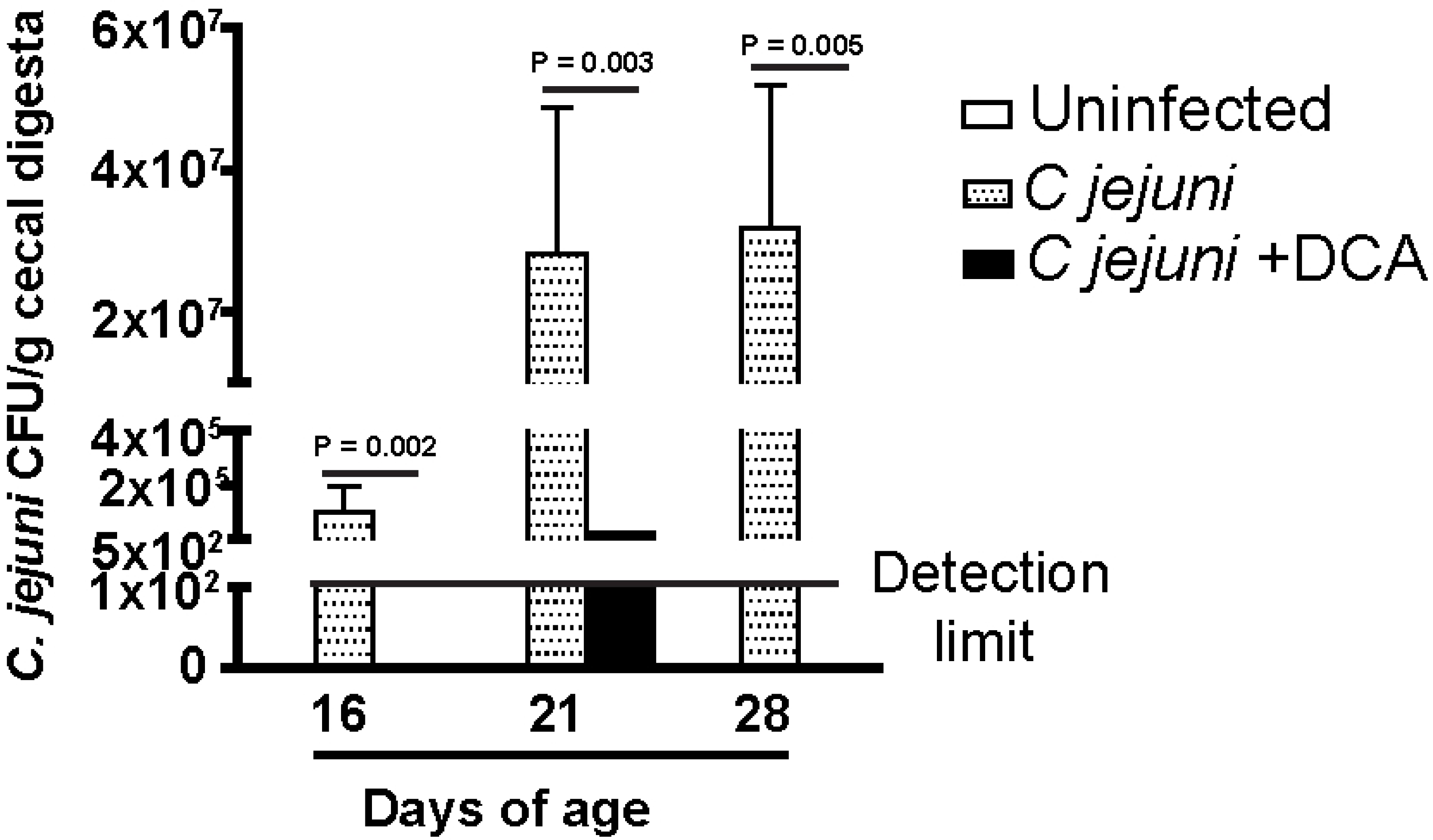
DCA prevents against *C. jejuni* stain 81-176 colonization in chickens. Cohorts of 13 one-day-old broiler chickens were fed 0 or 1.5 g/kg DCA diets. The birds were infected with *C. jejuni* 81-176 at 14 days of age. Cecal digesta were collected at 16, 21, and 28 days of age, serially diluted, and cultured on *C. jejuni* selective medium at 42 °C. *C. jejuni* was counted after 48 hours of culture. All graphs depict mean ± SEM. Significant if P<0.05. Results are representative of 2 independent experiments.

### DCA promotes bird growth performance

Increased level of secondary bile acids DCA has been associated with obesity [36], but the role of DCA on animal growth is unclear. To investigate the contribution of DCA on chicken growth, the bird growth performance of body weight gain was measured at 14 and 28 days of age with or without *C. jejuni* infection. Unlike the outcome of severe intestinal diseases when infecting to human or mice [43,44], *C. jejuni* infection in chickens neither induced diseases nor reduced the bird growth performance on body weight gain compared to uninfected birds (Figure 2A). Remarkably, DCA promoted growth performance of body weight gain by 36.3 % at 14 days of age, compared to control birds. The body weight gain of birds fed DCA diets posed 12.7% increase compared to infected control birds at 28 days of age (Figure 2B). Interestingly, *C. jejuni* colonization increased chicken body weight gain during d 14-28. These findings suggest that DCA promotes bird growth performance of body weight gain and *C. jejuni* colonization doesn’t induce adverse effect on bird health and growth.

**Figure 2.**
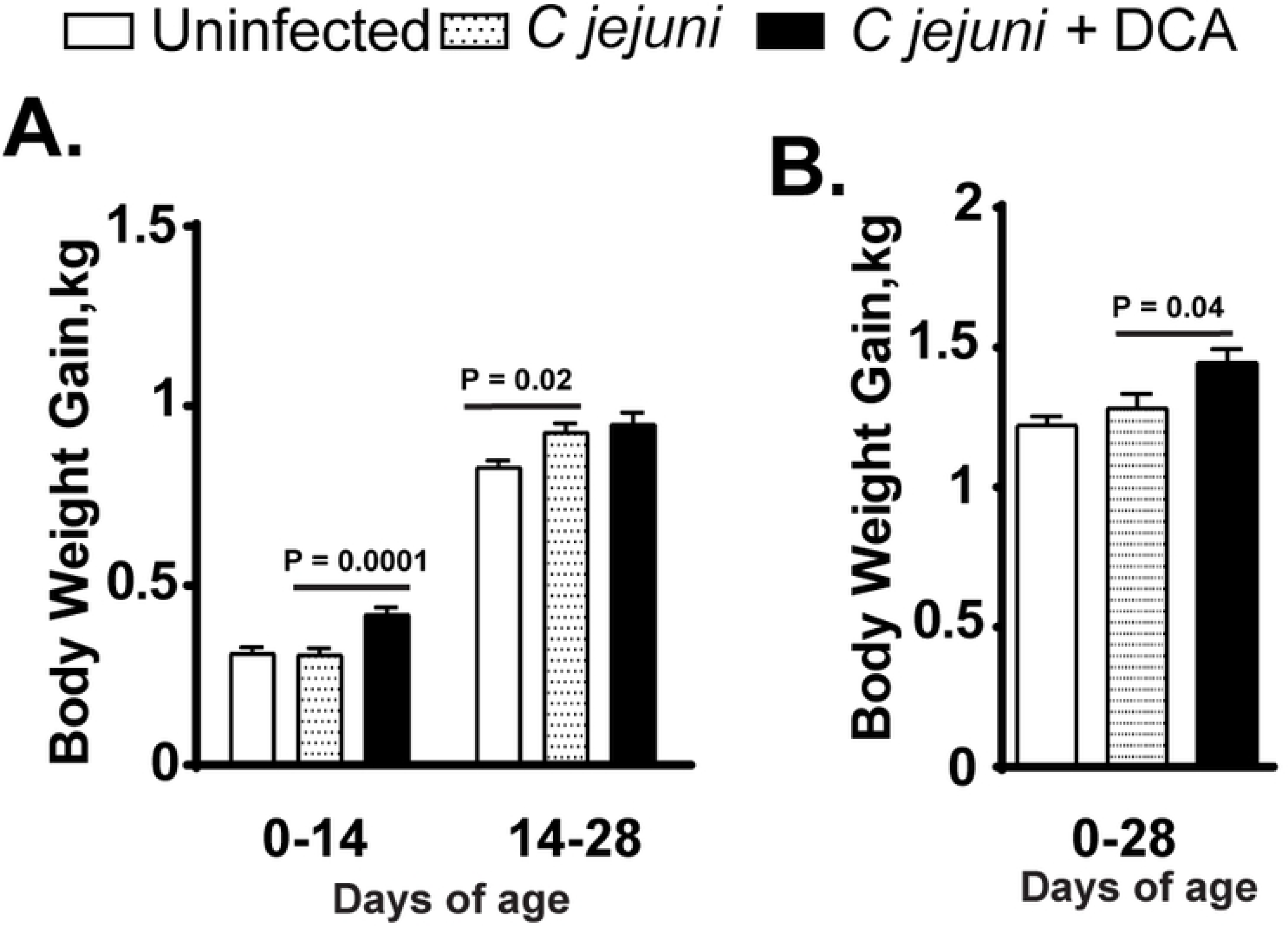
DCA promotes broiler chicken growth performance. Cohorts of 13 birds were fed DCA diet and infected with *C. jejuni* as described in Figure 1. (A) Periodic bird growth performance of body weight gain. (B) Accumulative body weight gain at 28 days of age. All graphs depict mean ± SEM. Significant if P<0.05. Results are representative of 2 independent experiments.

### DCA fails to inhibit *C. jejuni* 81-176 *in vitro* growth

Since DCA prevented *C. jejuni* colonization in chickens, we then examine whether DCA would directly inhibit *C. jejuni* 81-176 growth. The bacterium was inoculated in Campylobacter Enrichment Broth in the presence of different concentrations (0, 5, and 25 mM) of bile acids DCA, CA, and TCA and cultured at 42°C for 24 hours under microaerobic condition. *C. jejuni* growth in 5 and 25 mM DCA broth induced gel formation and the OD_600_ reading was not accurate, therefore plate counting was used. Notably, primary bile acid CA at 25 mM reduced *C. jejuni* 81-176 growth (Figure 3A). Interestingly, conjugated primary bile acid TCA and secondary bile acid DCA didn’t reduce *C. jejuni* 81-176 in vitro growth.

**Figure 3.**
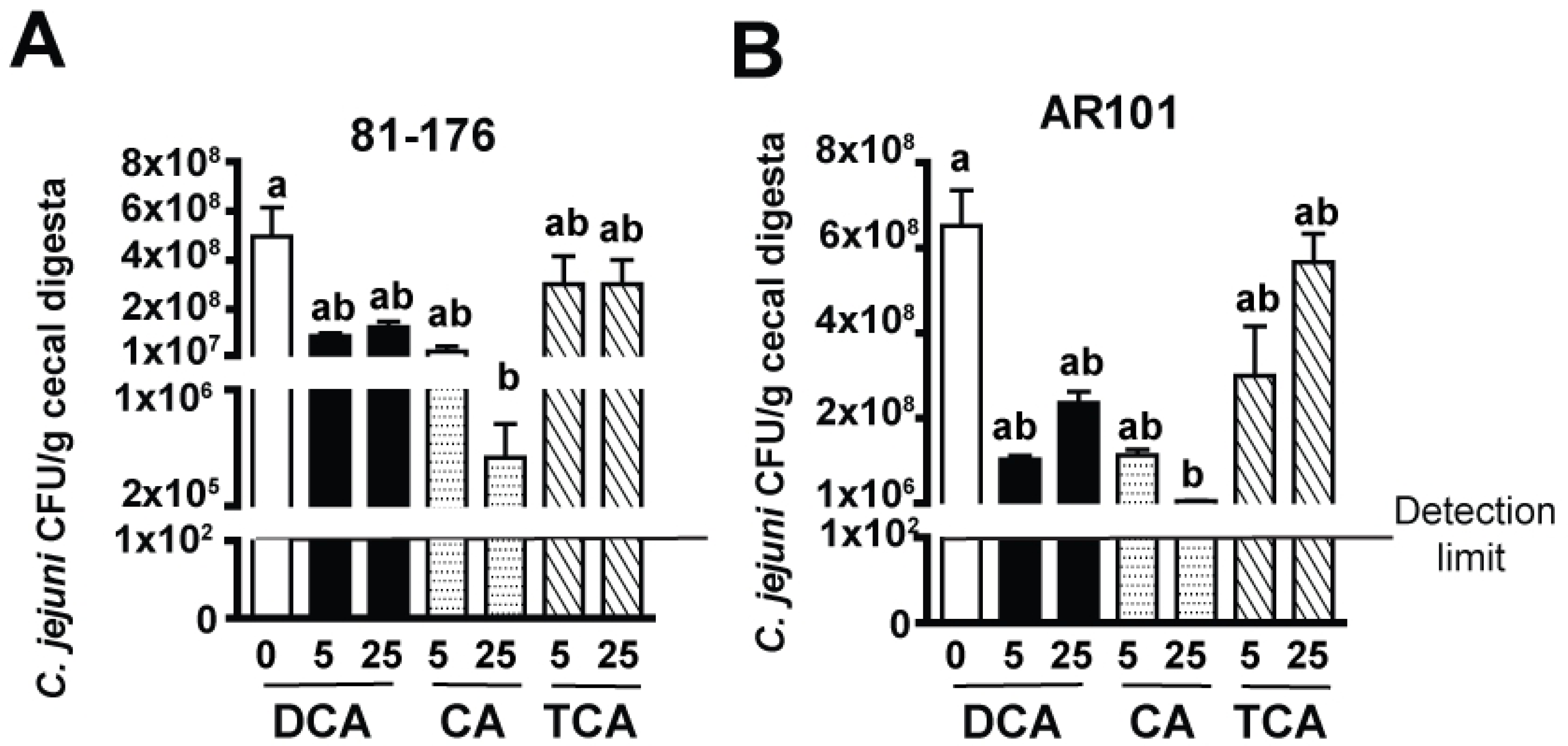
Bile acids fail to inhibit *C. jejuni in vitro* growth. *C. jejuni* 81-176 or AR101 at 10^3^ CFU was inoculated into 5 ml *Campylobacter* broth under microaerophilic condition and cultured at 42 °C for 24 hours. The broth was supplemented with various concentrations (0, 5, or 25 mM) of secondary bile acid DCA, primary bile acid CA, or conjugated primary bile acid TCA. (A) Bile acids on 81-176 growth. (B) Bile acids on AR101 growth. Different letters represent significance (P<0.05) between treatments. Results are representative of 3 independent experiments.

Our long-term goal is to use microbiome to reduce *C. jejuni* transmission in chickens and to attenuated transmitted campylobacteriosis using mouse model as reported before [39,44]. Hence, *C. jejuni* colonized and transmitted in chickens with microbiome treatment is required to be isolated for subsequently infecting *Il10^-/-^* mice. It was problematic when *C. jejuni* 81-176 couldn’t be isolated from DCA treated chickens as showed in Figure 1. Different *C. jejuni* strains show variable colonization ability in chickens [45]. Based on this knowledge, we then assessed the impact of bile acids on growth of *C. jejuni* strain AR101 which was isolated from chickens at Dr. Billy Hargis’s laboratory. Consistently, only 25 mM CA significantly reduced *C. jejuni* AR101 growth but not TCA or DCA (Figure 3B). Because DCA didn’t significantly reduce *C. jejuni* 81-176 or AR101 *in vitro* growth, it was concluded that DCA against *C. jejuni* chicken colonization results from factors other than DCA direct inhibition on *C. jejuni* growth.

### Other secondary bile acids don’t reduce *C. jejuni* AR101 colonization in chickens

Since DCA reduced *C. jejuni* 81-176 colonization in chickens, it was possible that other secondary bile acids also decreased the bacterial chicken colonization. To assess this possibility, birds were fed diets supplemented with 0 or 1.5 g/kg DCA, LCA, or UDCA from d 0. Because the chicken ages of *C. jejuni* AR101 infection didn’t influence its chicken colonization (Supplement Figure 1), birds were infected once with *C. jejuni* AR101 at d10 instead of d14 to have longer time to interact with the bile acids. Consistent with previous findings in Figure 1, DCA reduced 91% of *C. jejuni* AR101 colonization compared to infected control birds (2.06×10^6^ vs. 2.39×10^7^ CFU/g), while LCA and UDCA failed to decrease AR101 chicken colonization (2.05×10^7^ and 1.40×10^7^ CFU/g, respectively) (Figure 4). Together, these data suggest that only secondary bile acid DCA effectively reduces *C. jejuni* chicken colonization.

**Figure 4.**
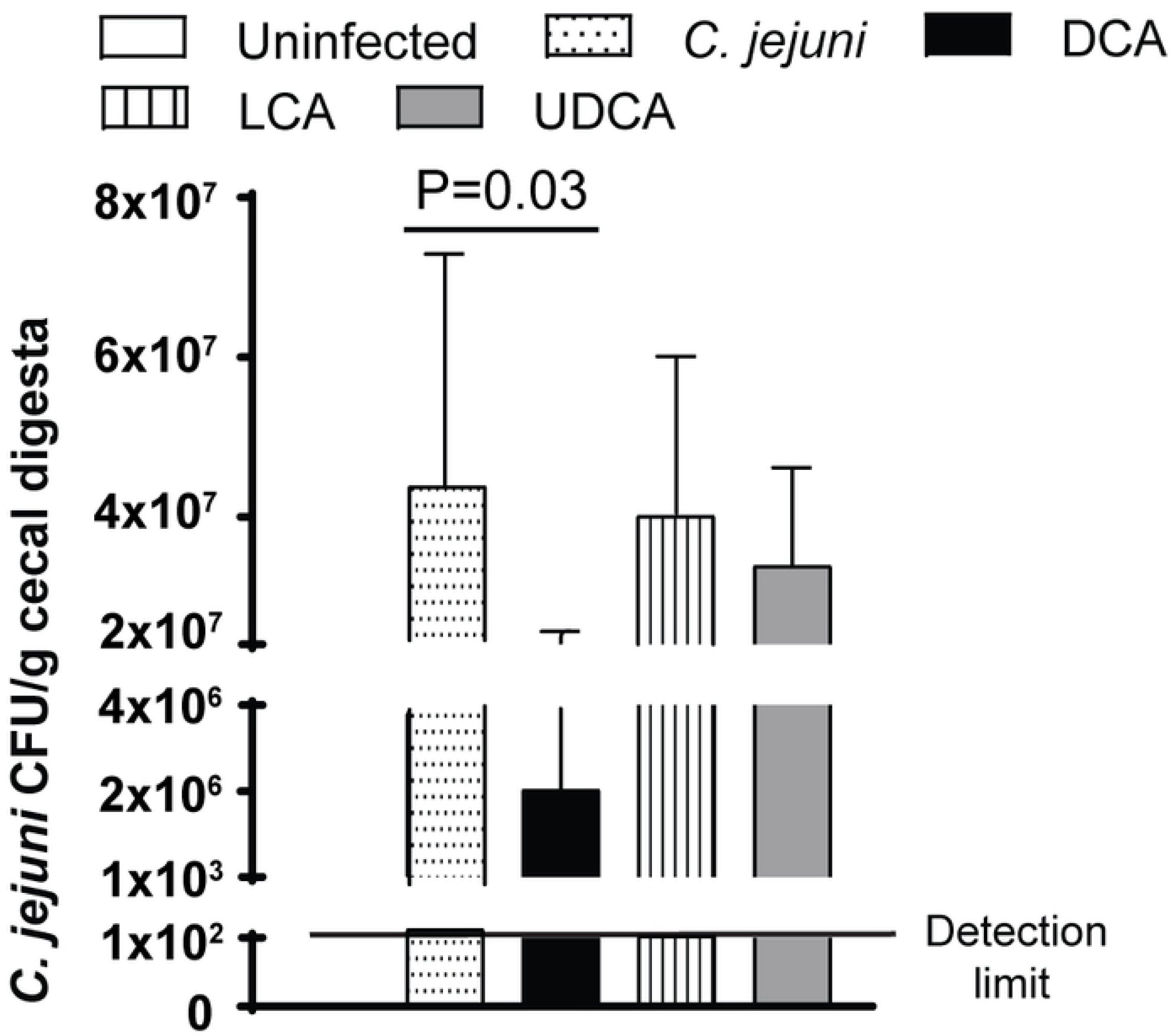
LCA and UDCA fail to prevent *C. jejuni* colonization in chickens. Cohorts of 18 chicks were fed diets supplemented with 0 or 1.5 g/kg of DCA, UDCA, or LCA. and were infected with *C. jejuni* AR 101 at day 10 of age. All birds were sacrificed and cecal digesta were collected at 28 days of age, serially diluted, and cultured on *C. jejuni* selective medium at 42 °C. *C. jejuni* was counted after 48 hours. All graphs depict mean ± SEM. Significant if P<0.05. Results are representative of 2 independent experiments.

### DCA modulates bird cecal microbiota

Relative abundance of phylum Firmicutes is dramatically expanded from 54 to 93% in intestine of rat fed primary bile acid CA [35]. Since DCA neither inhibited *C. jejuni in vitro* growth nor induced intestinal diseases (histopathology, data not shown) in chickens, we then reasoned that DCA modified microbiota against *C. jejuni* chicken colonization. To examine this hypothesis, cecal digesta from birds infected with *C. jejuni* 81-176 and fed DCA (Figure 1) were used to exact DNA. We selected these samples because *C. jejuni* wasn’t detected in the samples and the samples would be used for following experiment of isolating transplantation microbiota. Phylum specific primers were used to analyze the microbiota composition. Interestingly, *C. jejuni* infection didn’t change microbial composition in microbiota of infected vs. uninfected birds (Figure 5). Remarkably, DCA reduced phylum Firmicutes (82.7 vs. 98.8%) compared to infected control birds, while increased Bacteroidetes (16.9 vs. 0.8%). These results indicate that DCA is able to alter the chicken gut microbiota.

**Figure 5.**
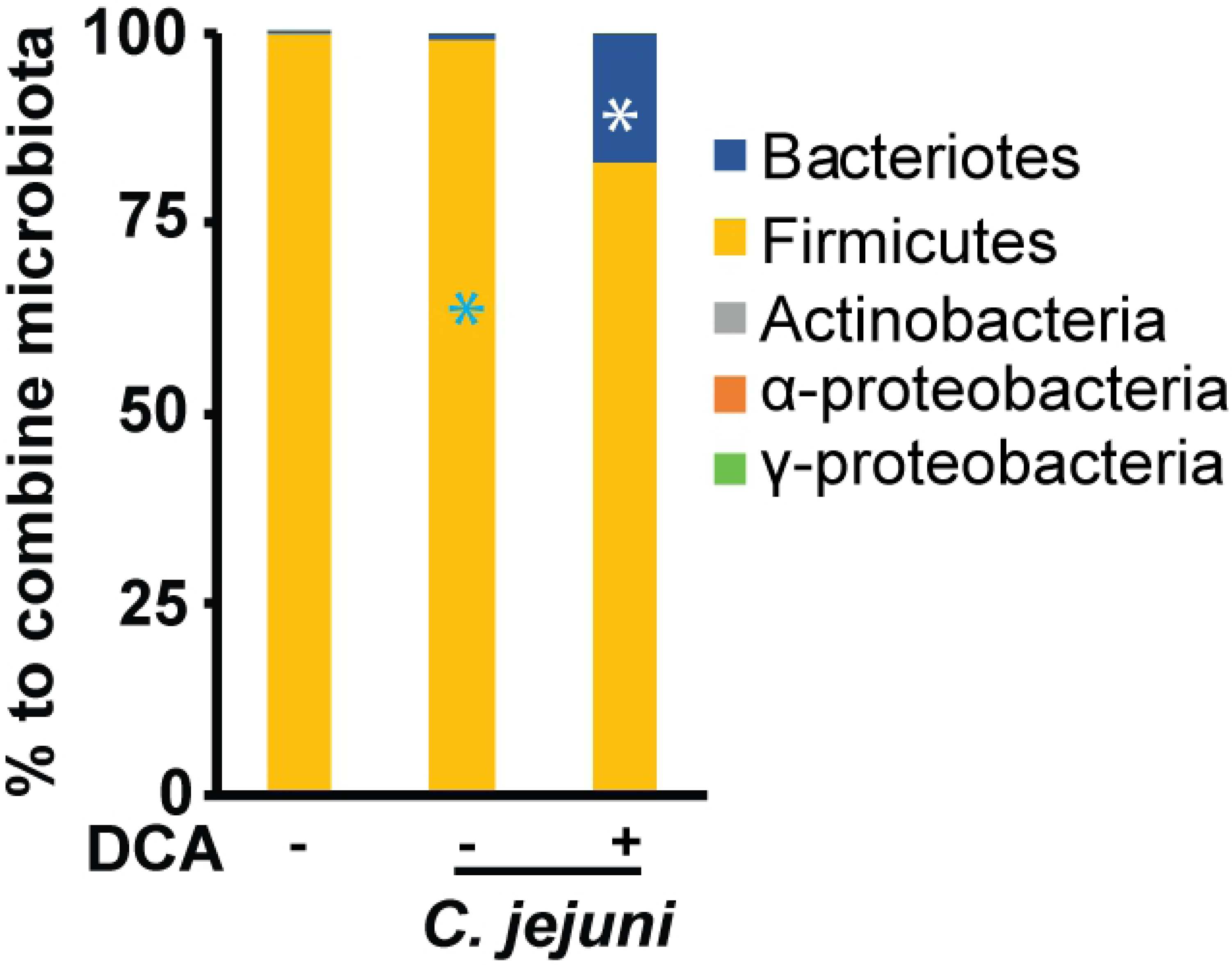
DCA modulates cecal microbiota. Cohorts of 13 chicks were fed 0 or 1.5 g/kg DCA diets and infected with *C. jejuni* 81-176 as described in Figure 1. Cecal digesta samples were collected and DNA was isolated. Real time PCR was performed to calculate bacterial composition at phylum level. All graphs depict mean ± SEM. *, P<0.05. Results are representative of 2 independent experiments.

### DCA modulated-anaerobes attenuates *C. jejuni* AR101 chicken colonization

Since DCA modulated chicken cecal microbiota, we then hypothesized that the altered microbiota contributed to the reduction of *C. jejuni* in chickens. In our previous studies, we found that only anaerobes prevent *C. jejuni*-induced intestinal inflammation in mice [39]. Based on this knowledge, cecal digesta from the birds fed DCA and infected with *C. jejuni* 81-176 in Figure 1 were used to culture bacteria on BHI plates under aerobic or anaerobic conditions. The resulted bacteria were labeled as DCA-modulated aerobes (DCA-Aero) or DCA-modulated anaerobes (DCA-Anaero), respectively. To functionally dissect the role of these newly isolated microbiota, birds were gavaged once with DCA-Aero or DCA-Anaero microbiota at 0 day of age. The birds were then infected with 10^9^ CFU/bird *C. jejuni* AR101 at 10 days of age.

Consistent with previous observations, *C. jejuni* colonization in chicken ceca reached a plateau at a level of 2.80×10^7^ CFU/g cecal digesta at 21 days of age (Figure 6A), or 11 days post infection. Importantly, DCA-Anaero significantly attenuated 93% of *C. jejuni* cecal colonization at 28 days of age compared to infected control birds (1.79×10^6^ vs. 2.52×10^7^ CFU/bird), while DCA-Aero only reduced 34.46% of *C. jejuni* (1.65×10^7^ vs. 2.52×10^7^) colonization. To examine if the transplanted microbiota colonized the chickens, cecal samples from the birds were collected. DNA was extracted and phylum level PCR was performed. Consistent with the inoculum DCA microbiota, the microbiota of DCA-Anaero birds showed increased Bacteroidetes (23.6 vs. 3.8%) but reduced Firmicutes (76.3 vs. 95.8%) compared to infected control birds (Figure 6B), while DCA-Aero birds shared similar microbiota composition to infected control birds. These data supported the notion of our successful DCA microbiota collection, inoculation, and chicken colonization. Taken together, these findings suggest that DCA modulates microbiota and the modulated anaerobes contribute to the reduction of *C. jejuni* colonization in chickens.

**Figure 6.**
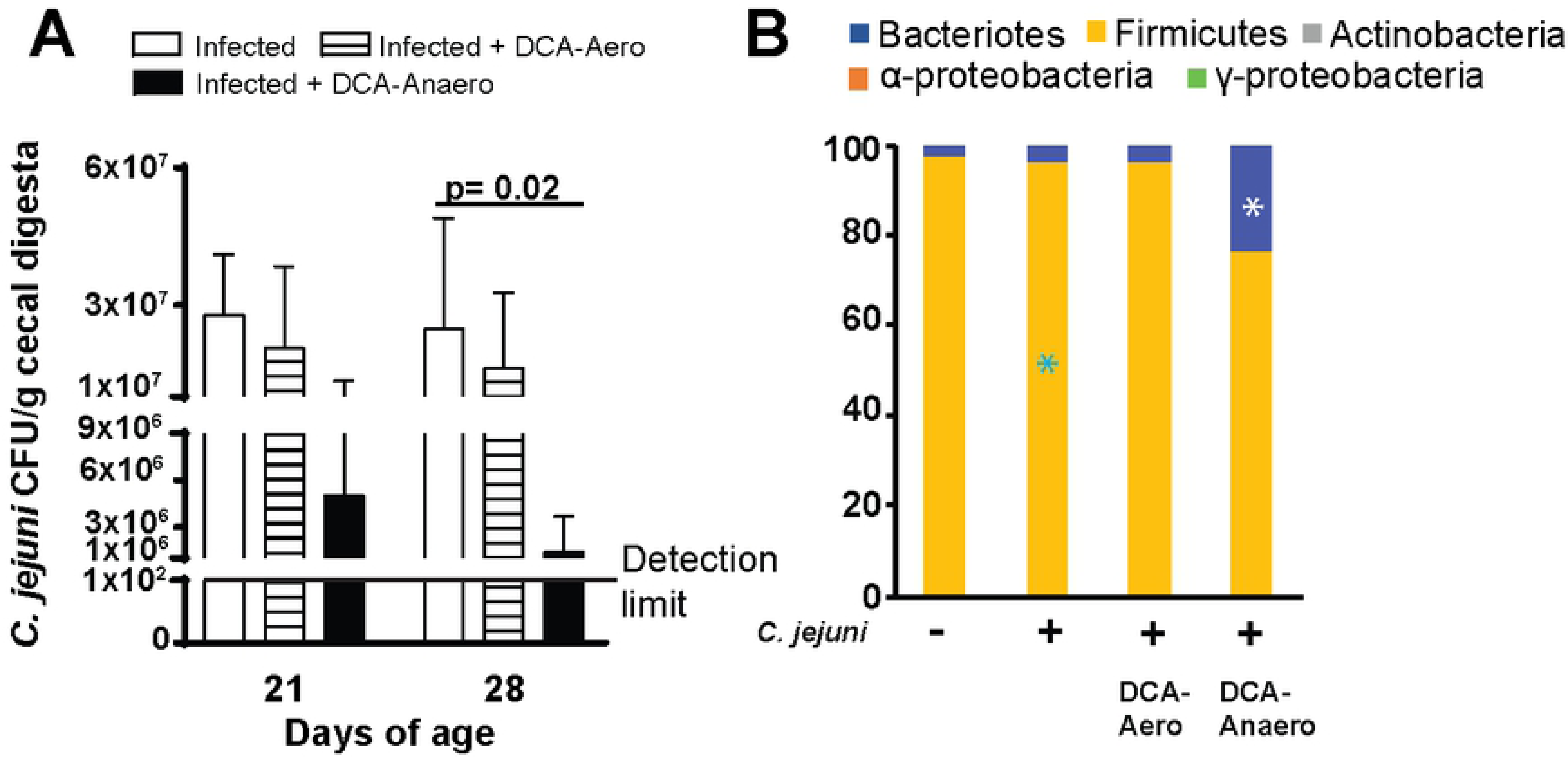
DCA-modulated microbiota attenuates *C. jejuni* colonization in chickens. Cohorts of 18 broiler chickens were orally transplanted with 10^8^ CFU/bird DCA-Aero or DCA-Anaero at d 0. The birds were then infected with 10^9^ CFU/bird *C. jejuni* AR 101 at 10 days of age. Cecal digesta were collected at 21 and 28 days of age, serially diluted, and cultured on *C. jejuni* selective medium at 42 °C. *C. jejuni* was counted after 48 hours. (A) *C. jejuni* cecal colonization. (B) Cecal microbiota composition at d28 at phylum level. *, P<0.05 between DCA-Anaero and infected control birds. All graphs depict mean ± SEM.; significant if P<0.05. Results are representative of 3 independent experiments.

## DISCUSSION

Although *C. jejuni* is one of the prevalent foodborne pathogens in developed countries, a paucity of information is available regarding reducing the pathogen in the main food animal source of chickens. Moreover, the microbiota and cellular events implicated in host resistance/susceptibility to *C. jejuni* infection remain elusive [46,47]. Here we report that microbial metabolic product DCA prevented colonization of both human clinical isolate *C. jejuni* 81-176 and chicken isolate AR101 in chickens. Interestingly, bile acids of DCA, TCA, or low concentration CA failed to reduce *C. jejuni in vitro* growth. Furthermore, neither bird ages of *C. jejuni* infection at d 5, 10 or 14, nor other bile acids of LCA or UDCA influenced the bacterial chicken colonization levels. Mechanistic studies revealed that DCA modified chicken cecal microbiota with increased phylum Bacteroidetes and reduced Firmicutes. Importantly, DCA-modulated anaerobes prevented *C. jejuni* chicken colonization. Altogether, these findings identified a novel mechanism that DCA shapes microbiota composition against *C. jejuni* colonization in chickens, suggesting a bidirectional interaction between microbiota and microbial metabolites.

A remarkable observation from our study is that DCA but not LCA or UDCA prevented *C. jejuni* colonization in chickens. The logic reasoning would be that the reduction would be through DCA directly impairing *C. jejuni* growth. DCA at 1.2 mM inhibits *C. jejuni* 81-176 *in vitro* growth after 12 hour incubation [48] but Lin and colleague found that the MICs of DCA and CA for *C. jejuni* 81-176 are 24 and 14 mM, respectively [49]. Interestingly, DCA at 48 mM fails to reduce *C. jejuni* 43431 growth at 6, 22, 25 and 30 hours of incubation but not at 16 hours [50]. Consistent with the latter reports, we found that conjugated primary bile acid TCA or secondary bile acid DCA at as high as 25 mM didn’t significantly inhibit *C. jejuni* 81-176 or AR101 *in vitro* growth. Consistent with Lin’s report [49], CA at 25 mM significantly reduced *C. jejuni in vitro* growth, suggesting a potential to manipulate this bile acid against *C. jejuni* chicken colonization. In animals, DCA reduces *C. jejuni* induced intestinal inflammation in ex-Germ Free mice without altering *C. jejuni* colonization level in colon [39]. Together, the knowledge and data suggest that DCA reduces *C. jejuni* colonization possibly through mechanisms other than directly inhibiting the bacterial growth.

Comprehensive database analysis showed that chicken microbiota at phylum level is mainly comprised of 13 phyla including Firmicutes (70%), Bacteroidetes (12.3%), Proteobacteria (9.3%), and other small proportion of Actinobacteria, Cyanobacteria, Spirochaetes, Synergisteles, Fusobacteria, Tenericutes, and Verrucomicrobia [51]. Our phylum level analysis of microbiota has found that birds fed DCA were colonized with reduced Firmicutes (82.7 vs. 98.8%) and increased Bacteroidetes (16.9 vs. 0.8%), which is associated with less *C. jejuni* colonization compared to control infected birds. Notably, this finding is consistent with a field survey report that birds from the farm with the highest *Campylobacter* counts are associated with the highest percentage of Firmicutes and the lowest percentage of Bacteroidetes, although microbiota composition is highly variable between inter- or intra-farms [52]. Interestingly, the microbiota in mice fed CA expands phylum Firmicutes (54 to 99%), class Clostridia (70 to 39%), and genus *Blautia* (8.3 to 55-62%), in the expense of phylum Bacteroidetes (30 to 0.39%) [35]. The levels of bile acids are associated with microbial community dynamics in the gut [32]. Bile acids directly inhibit gut microbes [33] and ileal bacterial overgrowth through FXR-induced antimicrobial peptides [34]. However, it remains elusive how various bile acid species differentially influence intestinal microbiota composition. In the future, we will investigate the microbiota composition using 16S rDNA sequencing in birds fed DCA, LCA, and UDCA.

Based on the results of DCA altering microbiota, we hypothesized that the microbiota played roles on protecting against *C. jejuni* chicken colonization. To functionally dissect the protection of the DCA-modulated microbiota, microbiota culture and chicken microbiota transplantation were performed. Indeed, DCA modulated anaerobes reduced *C. jejuni* chicken colonization, while DCA modulated aerobes failed to do so. Phylum level microbiota composition analysis by real time PCR strongly support the successful microbiota collection, inoculation and chicken colonization. Microbiota prevents against *C. jejuni* chicken colonization because *C. jejuni* colonizes at higher level in germ free or antibiotics pre-treated chickens compared to conventional birds [53]. Specific members of microbiota play an important role against *C. jejuni* induced diseases. Microbiota diversity and the relative abundances of genera Dorea and Coprococcus in family Lachnospiraceae were higher in healthy travelers compared to individuals suffered with campylobacteriosis [47]. Mouse microbiota with higher level of genera *Clostridium XI*, *Bifidobacterium*, and *Lactobacillus* is associated with resistance to *C. jejuni* induced colitis [39]. The three genera bacteria metabolize conjugated bile acids into secondary forms [21] which prevent campylobacteriosis in mice [39]. Interestingly, probiotics *Bifidobacterium longum* PCB133 and a xylo-oligosaccharide fail to reduce *C. jejuni* chicken colonization using plate enumeration [54]. We are processing the cecal samples to extract bacterial DNA to run 16S rDNA sequencing and we expect that we will find specific bacteria in DCA-modulated anaerobes responsible for protection against *C. jejuni* chicken colonization.

Finally, we found that *C. jejuni* colonization levels was independent on the ages of birds infected. It remains controversial whether chicken ages of inflection play any role on *C. jejuni* chicken colonization. *C. jejuni* is detected in 40% of broiler chicken flocks at 4 weeks of age and in 90% of flocks at 7 weeks of age [55]. This infection pattern could be attributed to age-related resistance or the necessary timing for *C. jejuni* transmission within house. Birds from parent breeders colonized with high level *C. jejuni* strain 99/308 are resistant to 99/308 only at 8 days of age but not at 1 or 22 days of age [56], suggesting specific and limited protection from the breeders. However, Han and colleagues showed that *C. jejuni* strains Lior6 and 0097 colonize less in birds of d 22 compared to d 1, after 14 days post infection [49]. Because bird microbiota starts to assemble after hatch, the older birds in some farms might develop microbiota resistant to *C. jejuni*, as in Han and colleagues’ report [49]. Similarly, germ free mice transferred to SPF housing for 14 days resist against *C. jejuni* induced colitis [39]. In those research reports showing no age difference on *C. jejuni* bird colonization, the resistant microbiota assembly in the birds might be blocked by strict biosecurity measurements or no available resistant microbiota in the environment.

In conclusion, the microbiota metabolite secondary bile acid DCA decreases *C. jejuni* counts and modulates microbiota composition in the chicken intestine. At mechanistic level, DCA-modulated anaerobic microbiota may be responsible for protecting against *C. jejuni* colonization in chickens. These findings identified a novel mechanism that DCA shapes microbiota composition against *C. jejuni* colonization in chickens, suggesting a bidirectional interaction between microbiota and microbial metabolites. Simultaneously reconstituting both microbiota and microbial metabolites may render better therapeutic effect against campylobacteriosis or the pathogen colonization.

## Abbreviations

DCA: Deoxycholic acid
CFU: colony forming unit
DCA-Anaero: DCA modulated anaerobes
DCA-Aero: DCA modulated aerobes

## Author Contribution

B. A. and X. S. designed the experiments and wrote the manuscript with input from co-authors of M.A., Y.M. K., Y. H., and B. H. B.A. performed animal and *in vitro* experiments and most analysis with the help of co-authors of X. S., H. W., M. A., A.A., and M. B.

---

**Supplemental Figure 1.**
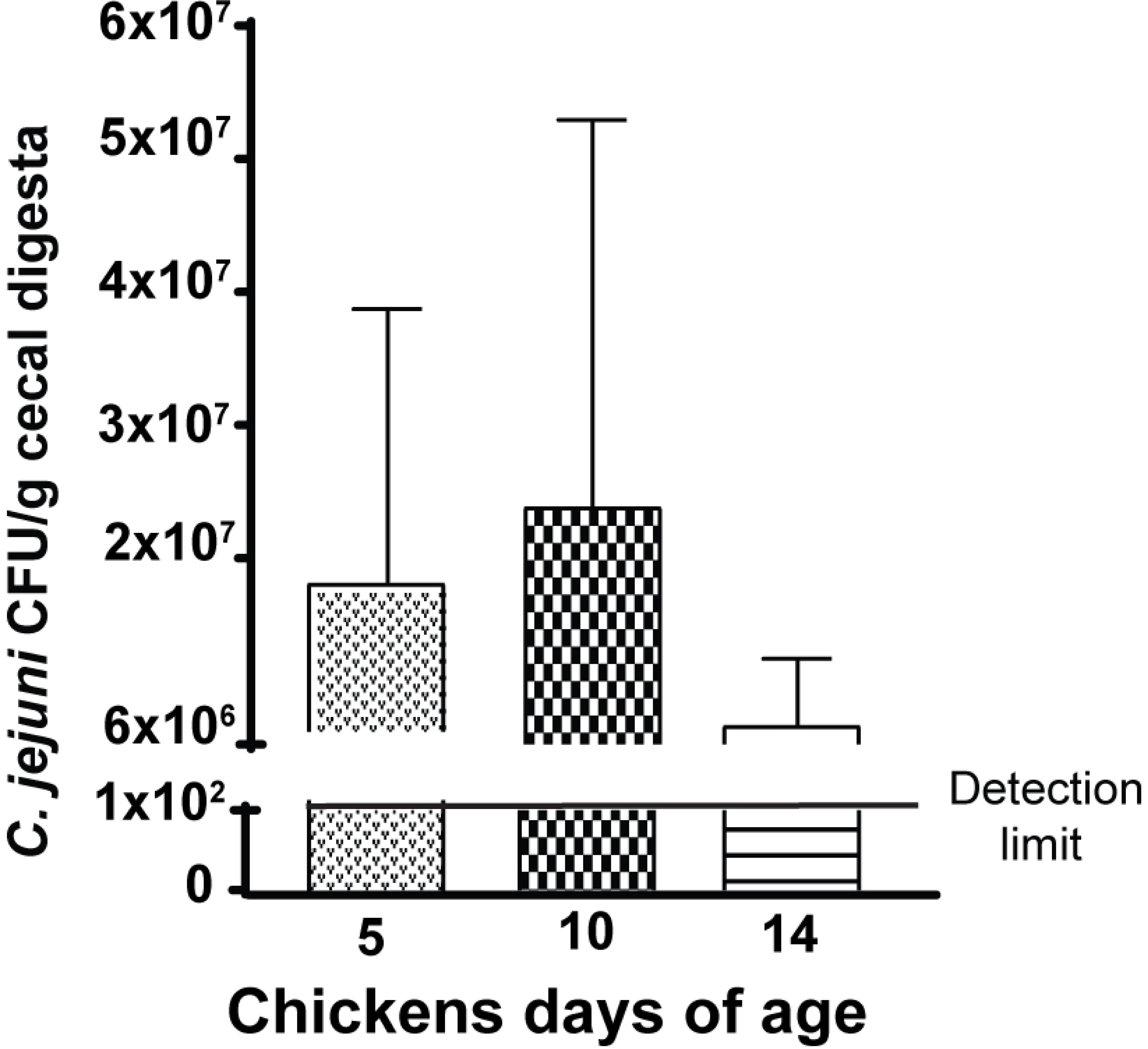
Chicken ages of infection don’t influence the bacterial colonization levels. Cohorts of 18 one-day-old broiler chicks were fed basal diet and orally gavaged with 10^9^ CFU of *C. jejuni* AR 101 at 5, 10, or 14 days of age. At d 28, birds were humanely sacrificed and cecal samples were collected. Cecal digesta samples were serially diluted and cultured on *Campylobacter* selective media. Colonies were enumerated and *C. jejuni* colonization levels were calculated. All graphs depict mean ± SEM. Results are representative of 2 independent experiments.

